# Hemin-binding DNA structures on the surface of bacteria promote extracellular electron transfer

**DOI:** 10.1101/2024.12.20.629652

**Authors:** Obinna M. Ajunwa, Gabriel Antonio S. Minero, Sissel D. Jensen, Rikke L. Meyer

## Abstract

Recent research has shown that bacteria in anoxic layers of *Pseudomonas aeruginosa* biofilms can respire by transferring electrons to oxygen via extracellular DNA (eDNA) and DNA-binding redox mediators that are unique to this species^1^. In this study, we propose a similar but generic mechanism by which bacteria can transfer electrons via DNA in biofilms, using hemin as a redox-mediator and hemin-binding G-quadruplex (G4) DNA structures in the extracellular matrix.

Using *Staphylococcus epidermidis* as a model organism, voltammetry showed that eDNA and hemin were needed for extracellular electron transfer (EET). Surface-associated G4-DNA formed a complex with hemin, which transferred electrons from the bacteria to an electrode under anoxic conditions. Addition of G4-DNA and hemin to growing biofilms promoted EET which was stable for days. G4-DNA/hemin is also a peroxidase-like DNAzyme, capable of transferring electrons from bacteria to H_2_O_2._

G4-DNA were only recently discovered to be abundant in the extracellular matrix of biofilms^2,3^. We now show that hemin turns these structures into conduits for EET. The study opens the door to new and generic mechanisms for bacterial energy conservation under oxygen-limiting conditions, and for tackling H_2_O_2_, a common host defense mechanism against bacterial infections.

## INTRODUCTION

Bacterial electroactivity describes a process by which bacteria use extracellular electron transfer (EET) to move electrons to or from metabolic processes in the cell^4,5^. This concept was previously thought to occur in a few prokaryotes that contain organelles or produce specific redox mediators^6^. However, a new report suggests that EET might be a ubiquitous phenomenon among microorganisms^7^.

EET is particularly beneficial for bacteria that use solid electron acceptors or that live in steep chemical gradients where soluble electron acceptors are out of reach. Biofilms create such gradients by consuming oxygen, and bacteria in the anoxic layers can benefit from using EET to transfer electrons to the oxic layer. In biofilms, bacteria are connected by a matrix of polymeric substances, such as polysaccharides, lipids, peptides, proteins and nucleic acids. Several of these are conductive and have been implicated in EET, e.g. polysaccharides in *Geobacter sulfurreducens*^8^ and polypeptides in *Bacillus subtilis*^9^. Recently, extracellular DNA (eDNA) was also implicated in EET. *Pseudomonas aeruginosa* transferred electrons from anoxic to oxic layers in biofilms via eDNA and small DNA-binding redox-active molecules called phenazines (pyocyanin and phenazine carboxamide). The soluble phenazines transferred electrons from intracellular metabolic processes to the extracellular “eDNA grid” and subsequently to oxygen^1^.

This mechanism for EET is unique to the pyocyanin-producing *P. aeruginosa*. However, DNA is a common biofilm component, and we therefore wondered if a similar but more generic principle for EET exists, where DNA-binding redox-active molecules transfer electrons from bacteria to a conductive eDNA network. We and others recently showed that eDNA and eRNA in biofilms can contain a variety of secondary structures: Z-DNA, i-motifs and G-quadruplexes (G4-DNA)^2,3,10,11^, and these are scattered throughout a net-like DNA superstructure in the biofilm’s extracellular matrix. The planar guanine quartets of G4-DNA have an increased potential for π-stacking and electron orbital sharing^12^, which leads it to form a stable complex with hemin – one of the most abundant redox mediators in biology. The G4-DNA/hemin complex is also a DNAzyme with oxidoreductase activity. We hypothesized that G4- DNA/hemin complexes can facilitate EET in biofilms, and that this is a generic mechanism for DNA-mediated EET.

The aim of this study was to provide proof-of-concept for G4-DNA/hemin-mediated EET in bacteria, and to demonstrate how these structures can boost EET in biofilms. We first determined if G4-DNA/hemin could interact with bacteria and facilitate EET. We then characterized the electrochemical signatures and charge transfer mediated by G4-DNA/hemin associated with both planktonic bacteria and biofilms of *S. epidermidis*. *S. epidermidis* is one of the most common culprits of implant-associated infections, and its main virulence factor is the ability to form biofilms. We used *S. epidermidis* as the model organism in this study due to its clinical relevance and because it was previously reported to produce large amounts of eDNA in the biofilm matrix – including G4-DNA structures^3^.

Our results support a scenario by which G4-DNA/hemin complexes associate with extracellular DNA and support bacterial metabolism through EET. With this knowledge, we believe that EET may provide a mechanism for bacteria in biofilms to sustain metabolic activity by long-range transfer of electrons to oxygen or hydrogen peroxide in the infectious microenvironment.

## RESULTS

### *S. epidermidis* can interact electrochemically with hemin via extracellular DNA (eDNA)

Our first aim was to establish if eDNA and hemin can confer electrochemical activity in *S. epidermidis* biofilms. We used voltammetry to compare the electrochemical signatures of biofilms with high and low amounts of eDNA. Biofilms of *S. epidermidis* 1457 wildtype or an eDNA-deficient strain (*S. epidermidis* 1457 Δ*atlE)* grew directly on the surface of electrodes for 48 h in media with hemin, and voltammograms (DPV and CV) confirmed a difference between the strains (Fig. 1 and extended Data Fig. 1). In CV analyses, the wildtype strain produced slight oxidation peaks at -0.10 V and +0.33 V and reduction peaks at -0.23 V and -0.37 V, but these peaks were not as distinct in the eDNA-deficient mutant strain (Extended Data Fig. 1).

**Fig. 1:**
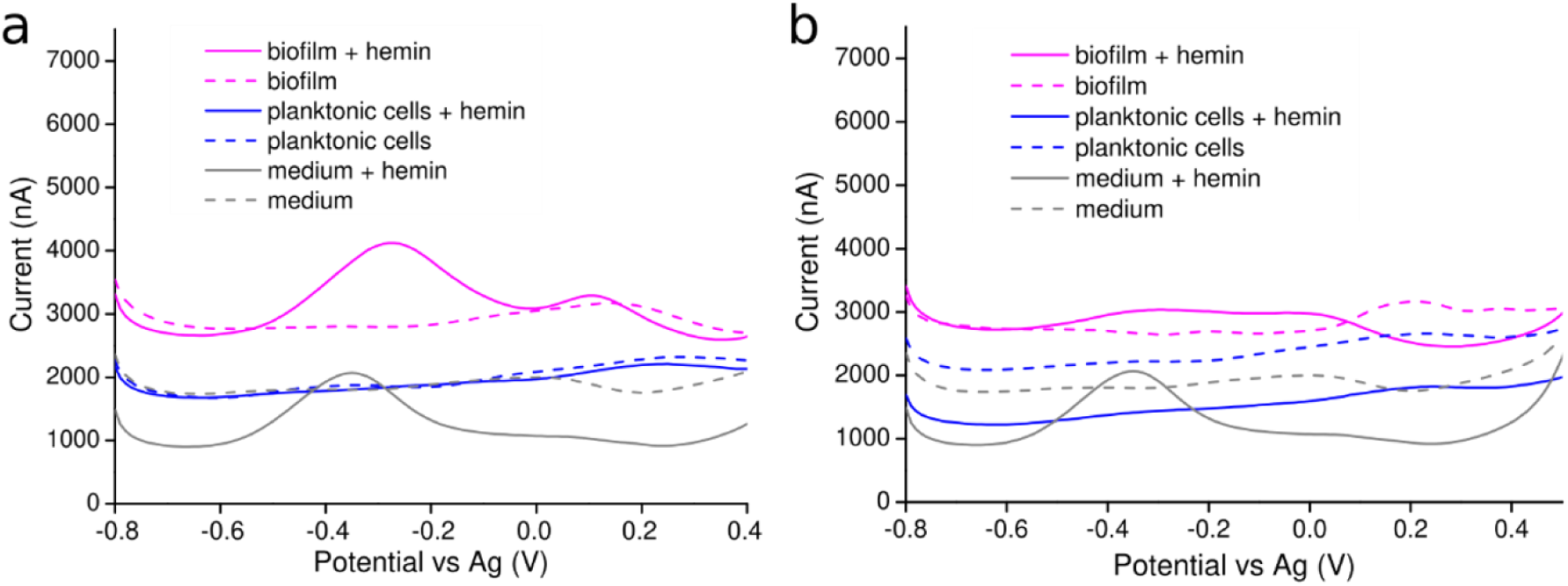
eDNA and hemin facilitate electrochemical signature of electroactive biofilms. Differential Pulse Voltammogram (DPV) of biofilms and planktonic cultures of **a)** *S. epidermidis* 1457 (eDNA producing) and **b)** *S. epidermidis* 1457 Δ*atlE* (non-eDNA producing) grown for 48 h in the presence or absence of 5 µM hemin. DPV curves are averages (n=3). Sterile medium (TSB, 0.2 M NaCl, +/- 5 µM hemin) served as control.

DPV analysis was more sensitive, and sterile controls contained a peak at -0.38 V resulting from dissolved hemin (Fig. 1a). Next, we grew planktonic *S. epidermidis* 1457 in media +/- hemin for 48 h and submerged the electrode to determine if planktonic bacteria could contribute to the electrochemical signal. These planktonic cultures contained no peaks – not even in samples with hemin (Fig. 1a), indicating that the bacteria removed hemin from solution, and that planktonic bacteria did not contribute to the electrochemical signature of biofilms in subsequent experiments.

In contrast to planktonic cultures, we observed two DPV peaks from biofilms grown on the electrode surface for 48 h in media with hemin (Fig. 1a). One peak (at +0.1-0.2 V) most likely represented the biomass, and this peak was independent of hemin in the media. In contrast, a very distinguished peak at -0.25 V was only present in biofilms grown with hemin, and we ascribed this peak to the electrochemical activity of hemin embedded in the biofilm. Importantly, the position of this peak was slightly shifted, relative to the peak from dissolved hemin, indicating that hemin interacts with another component - possibly eDNA.

To test this hypothesis, we conducted the same experiment on *S. epidermidis* 1457 Δ*atlE*, which forms biofilms that contain very little eDNA (Fig. 1b). The DPV peak associated with hemin in the biofilm was much smaller for this sample, which corroborates our hypothesis that hemin interacts with eDNA to confer electrochemical activity in biofilms.

### Hemin bound to G-quadruplex DNA can confer extracellular electron transport

We hypothesized that the electrochemical interaction between eDNA and hemin involves G4- DNA/hemin complexes. To test this hypothesis, we generated an experimental model in which *S. epidermidis* was loaded with G4-DNA on the surface, enabling us to study EET in bacteria with or without surface-associated G4-DNA/hemin.

Extracellular DNA co-localizes with exopolysaccharides in biofilms, and we therefore used *S. epidermidis* 1585 pTX*icaADBC*^13^ which has inducible production of poly-n-acetyl glucosamine (PNAG) to promote G4-DNA adsorption. We folded two different DNA oligos (c-Myc1 and c-Myc3) into G4-DNA (Extended Data Fig. 2), mixed them with *S. epidermidis*, and visualized surface-associated G4-DNA by fluorescence immunolabeling. We show that G4-DNA adsorbed primarily to PNAG-producing cells (Fig. 2a), and the G4 trimer c-Myc3 absorbed in larger quantity than c-Myc1 (Extended Data Fig. 4), resulting in higher electroactivity (Extended Data Fig. 5). We therefore chose c-Myc3 for our further analyses, and we used *S. epidermidis* 1585 pTX*icaADBC* +/- induction of PNAG production as positive and negative controls for bacteria +/- G4-DNA on the surface. *S. epidermidis* 1585 pTX*icaADBC* was used in all subsequent experiments unless specifically stated.

**Fig. 2.**
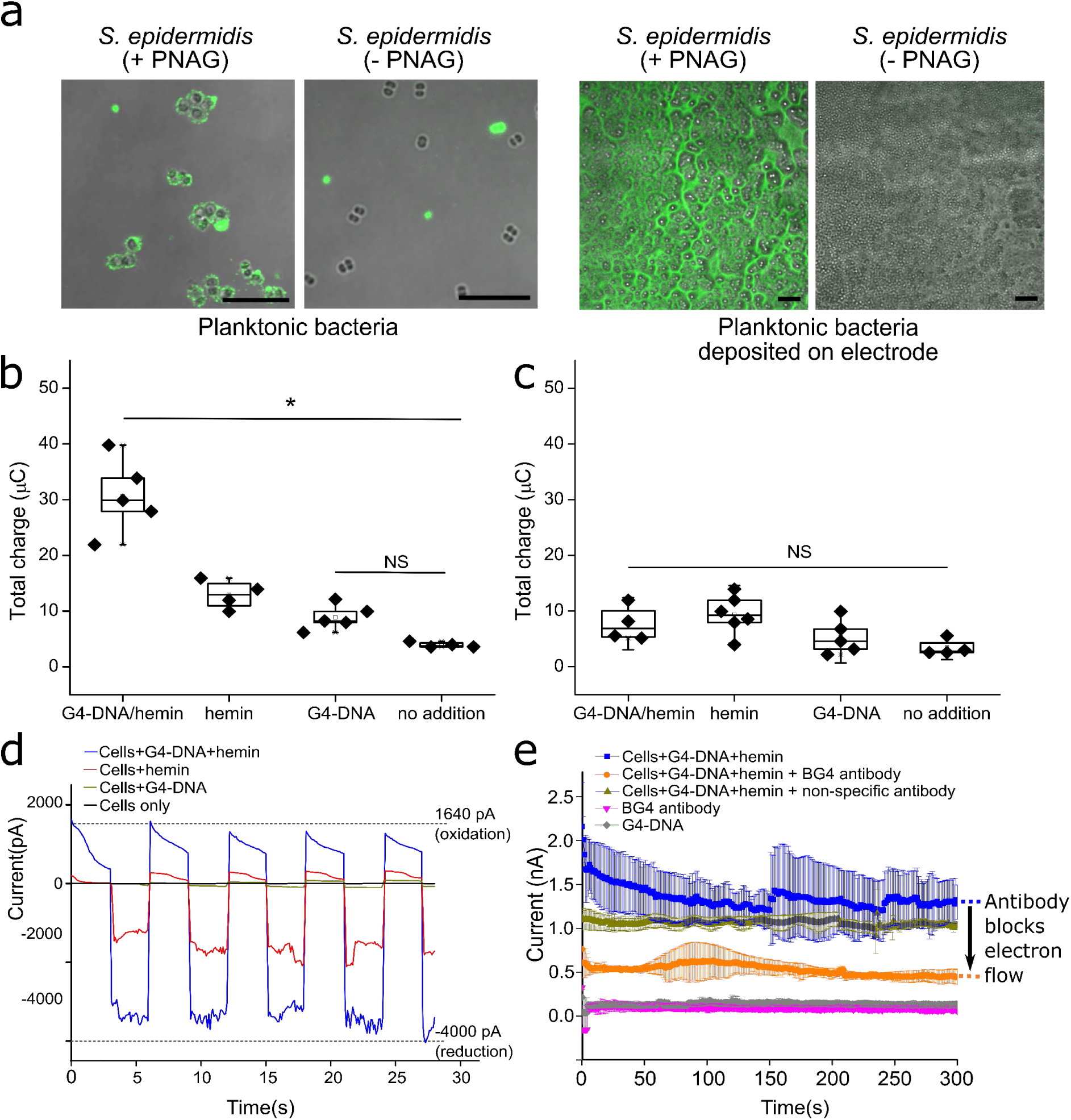
G4-DNA adsorbs on cell surface of PNAG-producing *S. epidermidis* and enables bidirectional electron flow. **a.** Externally added G4-DNA adsorbs to the surface of planktonic *S. epidermidis* if PNAG production is induced. Planktonic *S. epidermidis* were grown ±xylose for PNAG induction and incubated with G4-DNA for 1 h. The bacteria form a dense layer when deposited on an electrode for subsequent analysis (lower panel). Confocal laser scanning (CLSM) images show bacteria (brightfield) and G4-DNA (green) visualized by fluorescence immunolabeling with antibody BG4. Scalebar = 10 μm. **b.** Charge transfer from PNAG-induced *S. epidermidis*. Addition of G4-DNA/hemin generated significantly higher total charge after 300 s at 0.4 V poised potential. Hemin did not increase charge transfer if G4-DNA was absent. ∗ represents statistical significance (p < 0.05), ANOVA and Tukey’s test, NS= Not significant. **c.** Charge transfer from PNAG-deficient *S. epidermidis* that do not bind G4-DNA. Addition of G4-DNA and hemin did not affect the total electrical charge. **d.** G4-DNA/hemin on the surface of *S. epidermidis* could release and uptake electrons bidirectionally as demonstrated by short multi-step chronoamperometric analyses (MSCA). Values are an averages (n=3). Blocking the contact between G4-DNA/hemin and the electrode surface with G4-specific antibodies disrupts EET. A decline in the electron flow (indicated by black arrow) from planktonic *S. epidermidis* with G4- DNA/hemin was observed after interaction with G4-DNA antibody.

In an experiment to check that hemin was not toxic to the bacteria, we incubated *S. epidermidis* +/- induction of PNAG production and in the presence/absence of G4-DNA, and presence/absence of 5 µM hemin in the growth media. CFU enumeration of overnight cultures confirmed that hemin was not toxic, but it also revealed a 10-fold increase in CFU in samples with hemin. Notably, this increase in CFU only occurred in cultures with surface-adsorbed G4- DNA (Extended Data Fig. 3). Thus, hemin increased the growth yield when bacteria contained G4-DNA in the cell envelope.

Next, we grew planktonic cultures +/- induction of PNAG, added G4-DNA, hemin, or G4- DNA + hemin, and incubated for 1 h before washing and depositing the bacteria on the electrode surface for quantification of charge transfer at 0.4 V, which would capture the electrochemical potential of eDNA/hemin according to our results in Fig. 1. The charge measured in this experiment depends both on the number of bacteria on the electrode surface, and their ability to transfer electrons. We therefore visualized the adsorbed bacteria and G4- DNA within the adsorbed layer. Both strains adsorbed on the electrode surface as a dense layer, but only the PNAG-producing strain contained G4-DNA (Fig. 2a). Differences in charge transfer could therefore be ascribed to differences in mechanisms for electron transfer rather than differences in biomass.

The total electrical charge measured after short chronoamperometric analysis showed that EET activity was higher for bacteria with G4-DNA and hemin (Fig. 2b). In contrast, the total charge for bacteria incubated with G4-DNA alone was not significantly different from the control. Bacteria incubated with hemin showed a small increase in electrical charge compared to the control, but much less than bacteria incubated with both hemin and G4-DNA. This small charge may be attributed to direct electron transfer via hemin, or by hemin bound to G4-DNA structures naturally present on the bacterial surface. When repeating the experiment on *S. epidermidis* without PNAG induction and therefore unable to adsorb G4-DNA, none of the samples had a higher electrical charge than the control (Fig. 2c), supporting our conclusion that G4-DNA/hemin complexes facilitate electrical charge transfer.

### G4-DNA/hemin on the bacterial surface enables bi-directional electron flow

EET enables release of electrons from bacteria (electrogenicity) but could be used for electron harvesting (electrotrophy) as reported in electroactive prokaryotes involved in biocorrosion and element cycling^5,14^. The robustness of an EET mechanism can be shown in its ability to facilitate bi-directional electron flow, creating a tool for energy generation through electrical respiration^15^. We therefore sought to determine if G4-DNA/hemin in the bacterial cell envelope could convey bi-directional electron flow.

We evaluated the oxidative and reductive electron flow using short-span multi-step chronamperometry (MSCA) under oxidative (0.4 V) and reductive (-0.4 V) potentials, and quantified the current from *S. epidermidis* treated with G4-DNA, hemin, or both. We observed a larger difference in the current when poising the electrode for reduction (electrons flowing from the electrode) compared to oxidation (electrons flowing to the electrode) (Fig. 2d). The electron flow toward the electrode requires electron donors from bacterial metabolism, while electrons are acquired directly from the electrode surface when flowing in the opposite direction. The difference in current may therefore simply reflect the availability of electrons. The difference in reductive and oxidative current was only observed in samples supplemented with hemin, and it was amplified when both hemin and G4-DNA was present. This lends further support to our hypothesis that G4-DNA/hemin complexes promote electron flow.

### Electron flow requires direct contact between G4-DNA/hemin and the electrode

If the G4-DNA/hemin complex is specifically needed for electron transfer, we expect that G4- DNA/hemin must make direct contact with the electrode to facilitate electron flow. To investigate this, we coated the electrode surface with a G4-DNA specific antibody (BG4) prior to adsorption of bacteria with G4-DNA/hemin on the surface, in order to block the contact between G4-DNA/hemin and the electrode. We then monitored electron flow by chronoamperometry. As a control, we coated the electrode with a non-specific anti-mouse antibody. The non-specific antibody did not reduce electron flow (*p* value < 0.05), but the G4- DNA-specific antibody reduced the current by approximately 1 nA (Fig. 2e). We therefore conclude that G4-DNA/hemin must be in contact with the electrode to facilitate electron flow.

### G4-DNA/hemin incorporated into growing biofilms increases electron flow

After measuring charge transfer and EET from adsorbed planktonic bacteria, we investigated if G4-DNA and hemin could contribute to EET in biofilms where the G4-DNA integrate into a larger eDNA network that facilitates electron transfer over longer distances. We inoculated bacteria into media with G4-DNA, hemin or both, and grew biofilms directly on the electrode surface (poised at 0.4 V) under anoxic conditions, prompting the bacteria to use the electrode as electron acceptor. We measured the current produced throughout the 48 h of growth, and subsequently visualized the biofilms by microscopy.

As expected, addition of G4-DNA did not promote a current from the biofilm to the electrode, while addition of hemin did. The increase in current from hemin was, however, temporary and started to decrease after 15 h. In contrast, the combination of G4-DNA and hemin resulted in a stronger and more stable current throughout the 48 h incubation (Figure 3A). While the G4- DNA in this experient were not generated naturally in the biofilm, our data is proof-of-concept for the ability of G4-DNA structures to promote EET via hemin in biofilms.

**Fig. 3.**
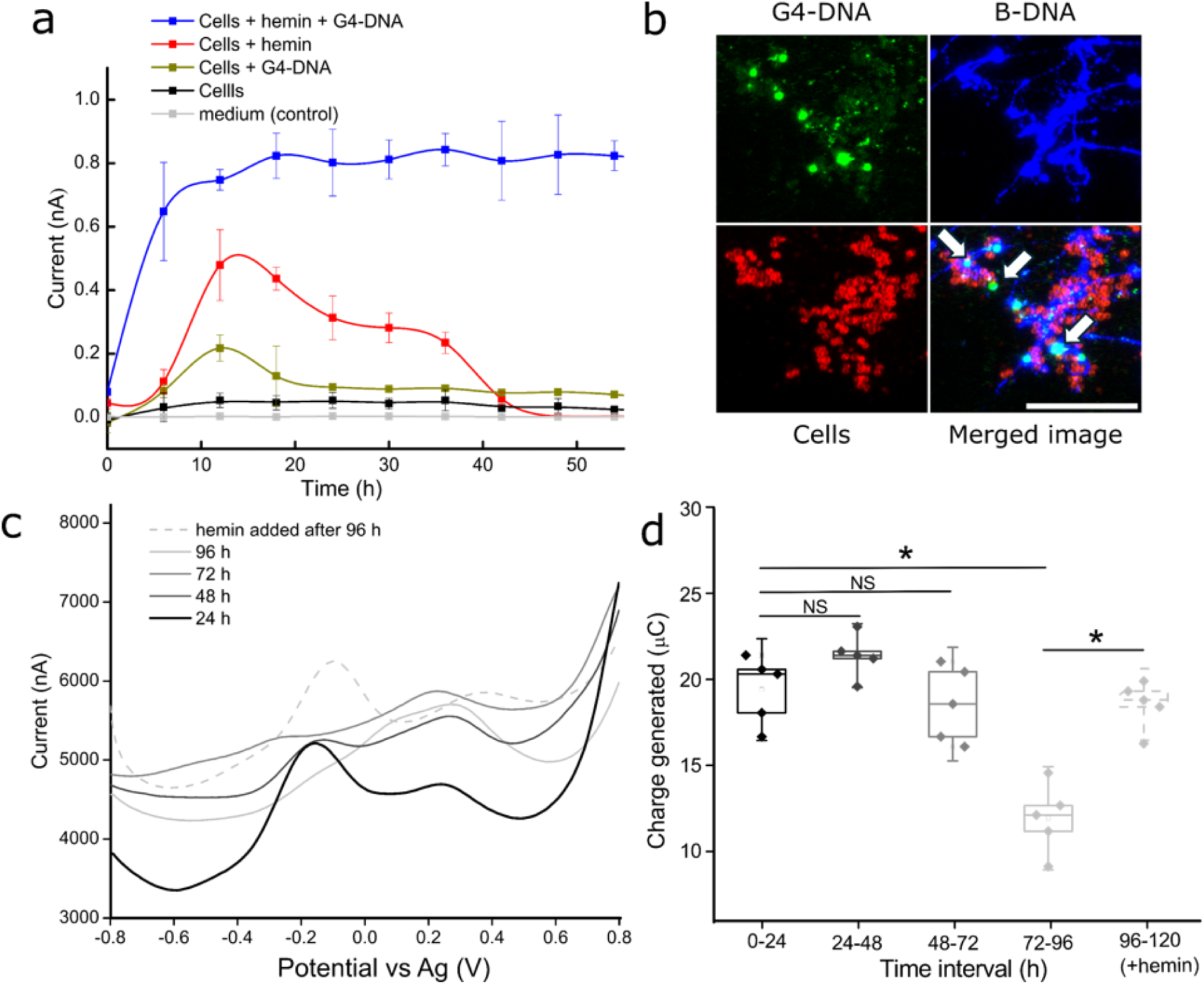
Biofilms grown with G4-DNA and hemin exhibit electroactivity for up to 48 h and sustain electroactivity for longer with hemin replenishment. **a.** Chronoamperometric measurements of *S. epidermidis* biofilm grown for 48 h on electrode surface poised at +0.4 V potential +/- addition of G4-DNA and hemin. **b.** CLSM images of *S. epidermidis* biofilms formed on electrode surface after 48 h growth with G4-DNA and hemin. G4-DNA forms nodular structures (indicated by white arrows) that associate with B-DNA in the biofilm. Scalebar = 20 µm. **c.** DPV of *S. epidermidis* biofilms grown with G4-DNA and hemin on electrode surface at +0.4 V potential. The current peak decreased after 24 h but could be recovered by replenishment of hemin. Charge generated under different times tested from 0 h to 96 h, showing decrease in electroactivity close to the 96 h age of the biofilm in comparison with 120 h old biofilm treated with additional hemin immediately after 96th hour (∗ represents significant difference between labelled experimental conditions (p < 0.05) based on ANOVA and Tukey’s test, NS: Not significant)

Visualization of biofilms confirmed that G4-DNA was incorporated into the eDNA matrix (Extended Data Fig. 6), and some G4-DNA formed nodule-like structures (Fig. 3b, Extended Data Fig. 8). Similar DNA structures have been reported with electroactive *P. aeruginosa* biofilms^1^, but they did not investigate if the biofilms contained G4-DNA. G4-DNA can aggregate through π-π interactions^12^, and they can also cause liquid-liquid phase separation in environments with molecular crowding^16^. It is not yet known how the nodule-like structures form and to which extent they contribute to electron transfer in the biofilm.

### EET stability requires replenishment of hemin

Our initial results indicated that EET in biofilms was stronger and more stable over time when G4-DNA/hemin was incorporated into the biofilm matrix (Fig. 3a). To test the boundaries for how stable EET is over a longer time frame, we incubated biofilms for up to 96 h in a similar experiment and carried out DPV analyses on the biofilms every 24 h. The DPV peak at approx.. + 0.1-0.3 V ascribed to biomass increased from 0-48 h incubation and then remained stable, indicating that biofilms growth stagnated after 48 h incubation but subsequently stayed associated with the electrode (Fig. 3c). Stagnation in biofilm growth was expected as the media was not replenished at any time, and electron donors for metabolism would become depleted. In accordance with the previous experiment, the peak around -0.1 to -0.3 V ascribed to G4- DNA/hemin increased in height during the first 48 h, but subsequently decreased and disappeared at 96 h incubation (Fig. 3c). The decrease in EET at 96 h was further corroborated by measurement of charge transfer (Fig. 3d).

The decrease in EET could be caused by a decrease in metabolic activity as nutrients were depleted from the media, or it could be caused by inactivation of the G4-DNA/hemin complex due to e.g. oxidative damage or loss of Fe^2+^. To test if it was the latter, we added 5 μM hemin to the biofilm to replenish hemin in the G4-DNA structures. This resulted in restoration of the G4-DNA/hemin DPV peak (Fig. 3c), indicating that EET activity of the G4-DNA/hemin complex requires continuous replenishment of hemin to retain optimal activity over longer timescales.

### Iron in hemin plays a key role in electroactivity of G4-DNA/hemin

Hemin consists of protoporphyrin IX (PPIX) coordinating a ferric iron ion (heme B)^17^. To validate the role of iron in EET observed in this study, we substituted hemin with the iron-deficient PPIX and compared electrochemical signatures based on DPV and charge generated. We grew biofilms on the electrode surface in media +/- G4-DNA and supplemented with either PPIX or hemin for 48 h before DPV analysis. The DPV peak ascribed to free hemin (approx. - 0.35 V) was only observed in cell-free controls, indicating that free hemin had less interaction with the biofilm-covered electrodes (Extended Data Fig. 7a). Biofilms grown with G4-DNA and PPIX had only one prominent peak at approx. + 0.1-0.3 V, which we ascribed to biomass. In contrast, biofilms grown with G4-DNA and hemin had two peaks: One ascribed to biomass and one at -0.1 to -0.3 V ascribed to the hemin/G4-DNA complex, indicating that iron is required for electroactivity. Furthermore, quantification of the charge generated from these biofilms (Extended Data Fig. 7b) showed that electroactivity required the presence of iron in hemin, and this result further corroborates iron’s importance EET via G4-DNA/hemin.

### G4-DNA/hemin in the cell envelope facilitates electron transfer to hydrogen peroxide

G4-DNA/hemin is a DNAzyme with peroxidase-like activity^18^, and we therefore hypothesized that it also facilitates transfer of electrons from bacteria to hydrogen peroxide, which then becomes an electron acceptor for bacterial metabolism via EET.

We first confirmed the peroxidase activity of G4-DNA/hemin in *S. epidermidis* biofilms using tyramide signal amplification (TSA). In this assay, the DNAzyme activates fluorescently labeled tyramide in a reaction with H_2_O_2_, leading to immobilization of fluorescent tyramide to near-by tyrosine. Biofilms amended with hemin showed a low level of extracellular peroxidase activity – probably arising from G4-DNA/hemin complexes naturally present in the biofilm. However, biofilms amended with both G4-DNA and hemin were strongly fluorescent, confirming that the immobilized G4-DNA/hemin complex was catalytically active (Fig. 4a). We then tested if the DNAzyme can transfer electrons from the bacteria’s metabolism to H_2_O_2_ by characterising the electrochemical reduction of H_2_O_2_ in suspensions of *S. epidermidis* using cyclic voltammetry (Fig. 4b). Addition of G4-DNA to the bacteria did not stimulate H_2_O_2_ reduction, while addition of hemin resulted in some H_2_O_2_ reduction (Extended Data Fig. 9). However, the combination of G4-DNA and hemin resulted in highest reduction of H_2_O_2_ (Fig. 4b) apparent by the smaller CV oxidation and reduction curves (Fig. 4b). This trend was similar to the electroactivity measured under the same conditions (Fig. 3a), indicating that G4- DNA/hemin functioned as an electron conduit from the bacteria, transferring electrons from bacteria to H_2_O_2_. Collectively the data shows that extracellular G4-DNA/hemin could expand the repertoire of electron acceptors for *S. epidermidis,* and it might even protect the bacteria from H_2_O_2_ – one of the key components in how the immune system fights pathogens^19^.

**Fig. 4.**
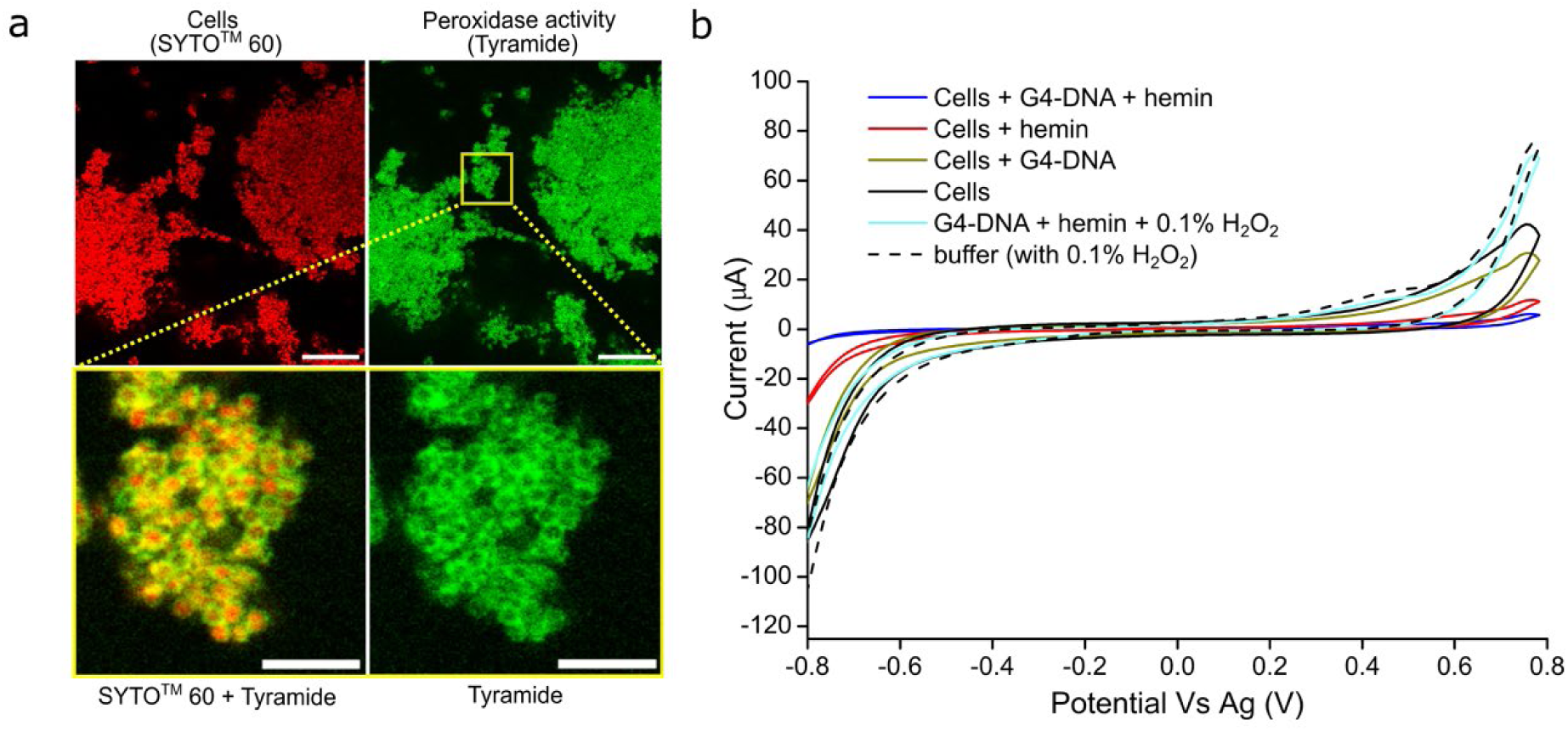
G4-DNA/hemin is an extracellular DNAzyme with peroxidase-like activity that could electrocatalytically degrade hydrogen peroxide. **a.** Tyramide signal amplification shows the location of extracellular peroxidase activity (green) in *S. epidermidis* (red, SYTO60^TM^ stain) grown with xylose for PNAG inducation and subsequent amendment of G4-DNA and hemin. Scale bar = 20 µm. Zoomed image shows that the catalytic activity is associated with the bacterial surface. Scale bar = 5µm. Electrocatalytic degradation of hydrogen peroxide with electrons from the bacteria. Cyclic voltammogram of electrochemical response from residual hydrogen peroxide in extracted peroxidase assay buffer after addition of cells, cells and hemin, cells and G4-DNA, or cells and G4-DNA/hemin complex.

## DISCUSSION

This study is the first to present a role for non-canonical DNA structures in microbial EET. We first established that EET in *S. epidermidis* biofilms requires the presence of eDNA. Since this species is known to form an eDNA network that contains G4-DNA, we hypothesized that the hemin-binding properties of G4-DNA played a role in DNA-mediated EET. The lack of charge transfer from soluble hemin in the absence of G4-DNA underlines the importance of the G4- DNA/hemin complex for EET (Fig. 2 b,c). G4-DNA might increase charge transfer by hemin simply by facilitating its proximity to the cell, but charge transfer could also increase due to enhanced electroactivity of hemin in the G4-DNA/hemin complex^20,21^ caused by the π-π stacking in G-quartets and the planar porphyrin structure of hemin^22^. Previous studies have shown that G4-DNA structures are an integral part of the eDNA network in staphylococcal biofilms^3^, and we propose a mechanistic model for EET in biofilms where hemin-binding G4- DNA enables electron transfer to and from an “eDNA grid” that collectively leads to flow of electrons through the biofilm matrix (Fig. 5).

**Fig. 5.**
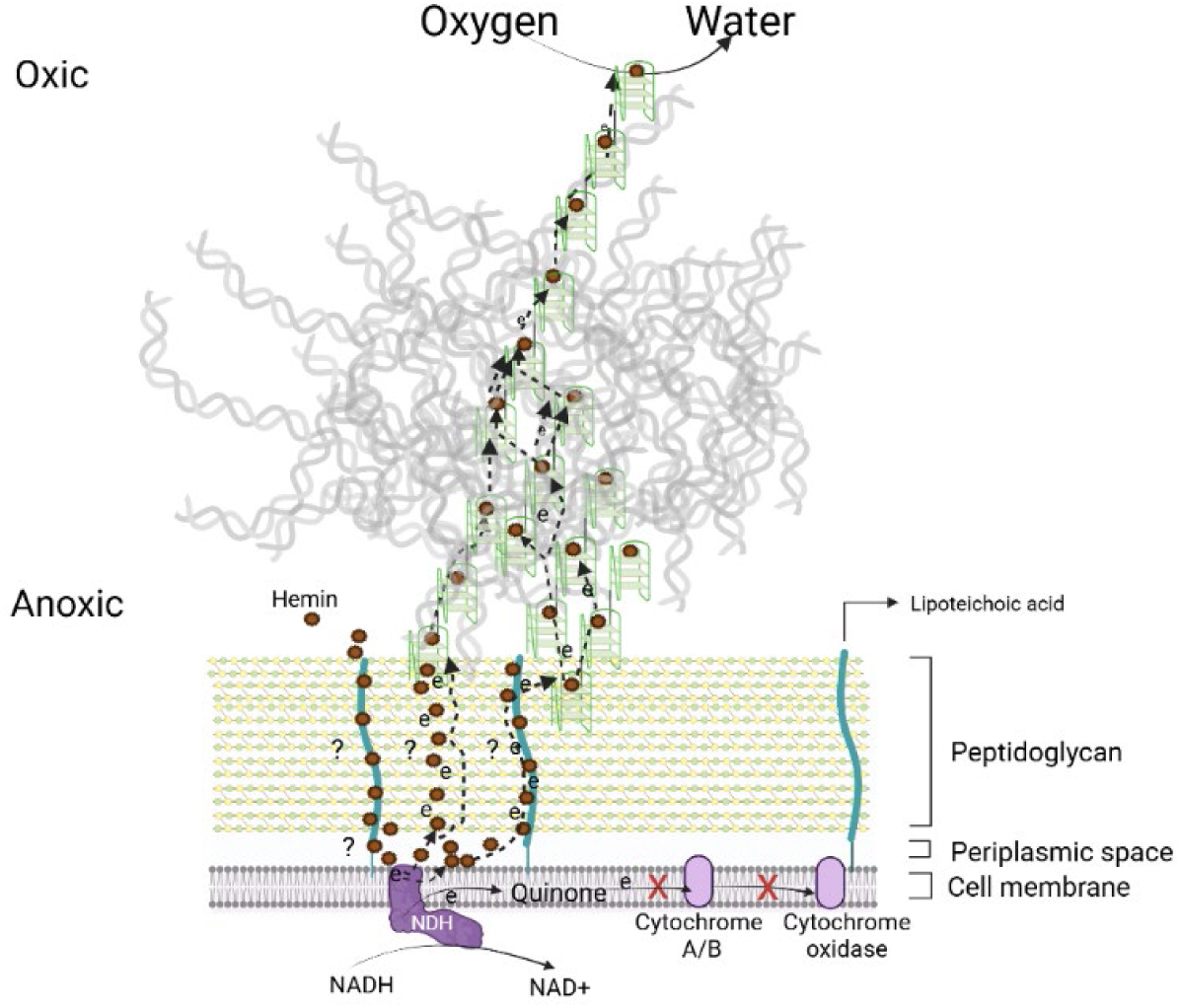
A conceptual model for the proposed involvement of G4-DNA/hemin in eDNA-mediated EET in biofilms. Created in BioRender. Ajunwa, O. (2024) https://BioRender.com/h06f025

Electrons from metabolic processes enter the electron transport chain (ETC) of *S. epidermidis* when NADH and FADH_2_ donate electrons to NADH dehydrogenase (NDH) or succinate dehydrogenase (SDH)^23^. NDH and SDH both transfer electrons to soluble menaquinone which transfers electrons to cytochromes and eventually to oxygen. In EET, electrons are diverted out of the cell. Based on the redox potential of hemin (-0.2 to -0.4 V vs Ag/AgCl), electrons are probably intercepted at NDH or SDH. NDH contributes to generating the proton motive force, and diverting electrons from NDH to hemin in stead of menaquinol will still result in ATP production.

How might the electrons be transferred from NDH on the surface of the cell membrane to G4- DNA/hemin outside of the cell wall? G4-DNA structures are too wide to pass through the 2 nm pores of peptidoglycan, and the bacteria must use another mechanism to move electrons up through the peptidoglycan layer, which spans tens of nanometers. Electron flow across the cell envelope differs in Gram positive and Gram negative bacteria. While many electroactive Gram negative bacteria use multiheme cytochromes^24^, these are uncommon in Gram positive bacteria such as *Staphylococcus*. In stead, Gram positive bacteria tend to secrete soluble redox mediators, such as nicotinaminde adenine dinucleotide (NAD) in *Bacillus subtilis*^25^, and flavins and quinones in *Bacillus cereus*, *Rhodococcus ruber*^26^ and *Lactiplantibacillus plantarum*^27^. *Listeria monocytogenes* immobilizes the secreted redox mediators in specific extracellular proteins (PplA) that span the peptidoglycan^28^. Another component that could facilitate electron transfer across the peptidoglycan are secreted conductive polymers. For example, *B. subtilis* forms long structures of poly γ-glutamic acid (γ-PGA) that make up a conductive biofilm matrix^9^.

In our case, hemin may perform a similar role as secreted redox mediators, but rather than being secreted, pathogenic bacteria can aquire hemin from blood. At the site of an infection, hemin enters the blood as a result of hemolysis or tissue damage, reaching levels of up to 25 µM^29^. While hemin binds to G4-DNA with high specificity, it can also interact with other biomolecules and thereby become immobilised in the cell wall through such interactions. Hemin possesses a hydrophobic ring with a positively charged iron core, enabling interaction with proteins^30^, lipids^31^, and membrane vesicles^32^. Due to its positive charge and planar geometry, hemin might interact with lipo- or wall-teichoic acids, which are negatiely charged and constitute up to 50% of the cell wall mass^33^. It is thus possible that hemin contributes to electron transfer across the peptidoglycan, either as a soluble or immobilised redox mediator.

### Comparison with DNA-mediated EET in other bacteria

DNA can be conductive in the right conformation and chemical environment. Canonical B-DNA allows electrical charge transfer between π orbitals of closely arranged base pairs^34^, and DNA structures are being considered for programmable nanoelectronics^35^ and batteries^36^. Notably, G4-DNA is more conductive than B-DNA because the stacked guanine tetrads facilitate π-π interactions^37^. Multiple G4-DNA can also stack into conductive G-wires reaching up to 100 nm^38^. This situates G4-DNA as a likely candidate for EET. DNA is the one matrix component that most biofilms have in common and G4-DNA structures have so far been reported in laboratory grown biofilms of *P. aeruginosa*^2^ and *S. epidermidis,* in implant-associated *S. aureus* infections^3^, and in dental plaque^39^. It is thus likely that G4-DNA is present in most biofilms, but its relevance was overlooked until now.

EET in biofilms can involve direct electron transfer via conductive biomolecules or indirect electron transfer using soluble redox-active molecules^4,5,15^ . Direct electron transfer via eDNA has been shown in *Shewanella*^40^, and a combination of direct and indirect electron transfer via eDNA and phenazines was reported for *P. aeruginosa*^1^. With the vast abundance of iron porphyrins in nature, we propose that G4-DNA and hemin provides a generic mechanism for EET that is conceptually similar to what was described for *P. aeruginosa*.

We show in this study that *S. epidermidis,* which is not percieved as an electroactive microorganism, does convey eDNA-dependent electron transfer in biofilms (Fig. 1). Moreover, EET could be stimulated and maintained when providing additional G4-DNA and hemin to the biofilms (Fig. 3a). We ascribe the stable EET to the immobilisation of hemin in the biofilm and integration of electroactive G4-DNA/hemin complexes in the wider network of eDNA. Furthermore, hemin in G4-DNA/hemin complexes may be better protected from oxidative damage compared to free hemin ^41^, and thus provide a more stable EET.

The use of DNA for EET requires a mechanism by which electrons can be transferred to and from the conductive DNA. While we propose that hemin is key to achieving such electron transfer, there are also other DNA-binding redox-active molecules, such as flavins and phenazines, which can increase DNA conductivity after being bound to DNA ^1,42^. Similarly chemosynthetic redox-active molecules such as methylene blue and monomethine cyanine dyes have also been demonstrated to bind to G4-DNA ^43,44^. The principle for eDNA-mediated EET described in this study may therefore go far beyond the specific mechanism described here.

### Biological implications of G4-DNA/hemin-mediated EET

In nature, EET is usually associated with bacteria that use insoluble electron acceptors. However, bacteria located in environments with steep chemical gradients can also use EET to reach soluble electron donors at some distance from the cell^1^. This ability enables continued respiration in anoxic microenvironments. In the infectious microenvironment, *S. epidermidis* biofilms most likely experience such steep oxygen gradients, making a case for how EET supports metabolic activity and survival of the population. The principle presented here is a generic mechanism through which biofilm-producing bacteria can sustain respiration in anoxic zones of a biofilm. As new research show that extracellular G4-DNA structures are common in biofilms, we raise the question of whether EET might also be more widely used by microorganisms than it was previously assumed.

Finally, we also show that H_2_O_2_ could be used as electron acceptor in EET due to the peroxidase activity of the G4-DNA/hemin complex. *In vivo*, host cells produce H_2_O_2_ in response to the infection^19^, and bacteria in the infectious microenvironment might therefore use H_2_O_2_ as an electron acceptor for respiration. In addition to facilitating respiration, the reaction may provide an additional benefit by removing H_2_O_2_, which is part of the host defense system^45^.

Another host defense mechanism is to limit the access of bacteria to iron^46^. Pathogens have therefore developed mechanisms to obtain iron from heme or hemin. For example, *S. aureus* contains near iron transporter (NEAT) domain proteins and Isd (iron-regulated surface determinant) for uptake of entire heme molecules from hemoproteins such as hemoglobin^47,48^. Furthermore, many staphylococci produce staphyloferrin siderophores that bind and assist the uptake of free iron. *S. epidermidis* lacks the machinery for heme uptake, but utilises sideophores such as staphyloferrin A^49^. Immobilisation of hemin on the cell envelope may serve as a mechanism for accessing iron in the infectious microenvrionment, and this phenomenon could explain the increase in growth yield caused by hemin when bacteria contained surface-associated G4-DNA (Extended Data Fig. 3).

## Acknowledgments

This work was funded by the Danish council for independent research, grant no. 2032-00294B. The authors thank Professors Lars Peter Nielsen and Andreas Schramm and the Center for Electromicrobiology (CEM) Aarhus University, Denmark for insightful discussions during the project and for access to CEM laboratories. We acknowledge financial support from the Danish National Research Foundation (DNRF136). The authors also thank Professor Dr. Holger Rohde of Universitätsklinikum Hamburg-Eppendorf, Germany, for providing us with the *Staphylococcus epidermidis* strains used in this work. Graduate student Elena Bouvier is also appreciated for assistance in preliminary laboratory work during her internship at the Interdisciplinary Nanoscience Center, Aarhus University, Denmark.

## Author contributions

**OMA** Investigation, methodology, data curation, data analyses, writing – original draft, writing – review and editing

**GASM** Methodology, writing – review and editing

**SDJ** Investigation, data curation, writing – review and editing,

**RLM** Fund acquisition, methodology, data curation, writing – original draft, writing – review and editing

## METHODS

### Chemicals and fluorescent stains

All chemicals used were first made as stock solutions. Hemin (Merck®) and protoporphyrin IX (Merck®) were prepared as 10 mM stock solutions in 200 mM Tris, 30 % DMSO and 100 mM NaOH (Merck®) at pH 11, and protected from light. Tris (Tris hydroxymethyl amino methane, pH 7.5, Merck®), modified MES buffer (25 mM MES (Sigma, M3671), 0.2 M NaCl (Sigma, S5886), and 0.010 M KCl (Sigma, 1.04936)) and DMSO stocks were prepared with filter-sterilized deionized water. Salts and sugars; NaCl, KCl and xylose were prepared as sterile filtered 1M stocks for buffer and culture preparations. Acetic acid (30% v/v) was used for pH adjustment of Tris buffers when prepared. Annealing buffer for G4-DNA was prepared by mixing 10 mM Tris, with 100 mM KCl (pH 7.2) and adjusting to pH 6.5 with acetic acid. Tyramide reagent was prepared by adding tyramide conjugated with Alexa Fluor 488 (1:100) (Invitrogen, B40953), 2 mM ATP (Thermo Scientific, R0441), and 0.1% hydrogen peroxide (Sigma, 216763) in modified MES buffer.

DNA binding stain SYTO^TM^ 60 (20 µM working concentration), was prepared as 100 x aliquots in sterile deionized water before use. A lipophilic dye FM 4-64 was prepared by mixing with sterile distilled water to 1mg/mL stock concentration. Working concentration for FM-464 was 10 µg/mL and aliquots for stains were made before storage at -20°C.

### Synthetic DNA, annealing and characterisation conditions

Oligonucleotides of guanine quadruplex (G4-DNA) sequences were purchased from Integrated DNA technologies (IDT), Belgium, as 100 µM stock in 1 mM Tris EDTA buffer. We used a monomer and a trimer of the c-Myc oncogene known to form G-quadruplex structures: c- Myc1: 5’- GA**<u>GGGTGGGTAGGGTGGG</u>**CGTCAACAGACTCGA-3’) and c-Myc3: 5’- GA**<u>GGGTGGGTAGGGTGGGGAGGGTGGGTAGGGTGGGGAGGGTGGGTAGGG TGG</u>**GCGTCAACAGACTCGA -3’. The part of the sequence that forms G-quadruplex is highlighted in bold. DNA annealing of oligonucleotides was done in an annealing buffer with composition stated above. Annealing was done by preparing 100 µL of 50 µM oligonucleotide stock in annealing buffer (pH 6.5), heating to 90°C for 3 min, and gradual cooling to 30°C. The structure of annealed oligonucleotides was confirmed by recording the circular dichroism (CD) spectrum using a J-810 spectropolarimeter at 220 – 320 nm, scanning speed 50 nm/min, and 2 nm bandwidth, and ultraviolet/visible spectrophotometer (UV/vis) also at 220 – 320 nm. For CD and UV/vis measurements, 5 µM oligonucleotides were prepared from sub-stocks and 70 µL sample was loaded into the spectropolarimeter and spectrophotometer respectively. CD signals were converted to ellipticity Δε (M^-1^ cm^-1^) using Beer-Lambert’s law and recorded.

### Strains, cultures and media

Tryptic soy broth medium (TSB, Merck) and Brain heart infusion medium (BHI, Merck) were prepared according to manufacturers’ specifications and used for culture preparations. Solid agar plates were prepared by mixing broth medium with 15 g/L agar powder. Autoclaved media broths supplemented with additional 200 mM NaCl were used for growing bacteria in overnight cultures. Bacterial cultures were made as overnight cultures by inoculating from single colonies on agar into 20 mL BHI/NaCl or TSB/NaCl broth depending on culture requirement and incubated for 15 – 20 h at 37°C with constant agitation at 150 rpm. *S. epidermidis* 1457, *S. epidermidis* 1457 Δ*atlE*, *S. epidermidis* 1585 pTX*icaADBC* were used in this study ^13,50–52^. Strains were stored at -80°C as stock cultures in cryotubes in 30% glycerol. For experiments with hemin, TSB/NaCl medium was supplemented with 5 µM hemin unless stated otherwise.

### Visualization of G4-DNA structures by immunolabeling

Fluorescence-conjugated antibodies were used for immunolabeling of DNA structures. G4- DNA specific antibody BG4 (goat monoclonal IgG lambda, Ab00174-24.1) with 0.02 % proclin was purchased as 1 mg/mL preparations in phosphate buffered saline (PBS) from Absolute Antibodies® with the fluorescent label Atto 488 and stored at 4°C. B-DNA specific antibody AB1.227156 (mouse monoclonal, AB27156, 1068371-2) was purchased from Abcam® without a fluorescent label. An anti-mouse antibody (goat monoclonal IgG H + L, XF346503 Invitrogen®) labelled with AlexaFluo405 was purchased as 2 mg/mL preparation in PBS with 0.05 % sodium azide and 50% glycerol and stored at -20°C and used as a secondary antibody for B-DNA detection (AB2). Both antibodies were mixed with 3 % bovine serum albumin (BSA) in PBS, which served as a blocking agent for non-specific binding during immunolabeling experiments.

Planktonic cultures were suspended for 2 min in 3% BSA in PBS before addition of BG4 antibody (working concentration 1:100 in 3 % BSA in PBS) and incubation for 90 min at room temperature. Immunolabeled bacteria were then briefly washed in 100 mM KCl and transferred to glass slides for microscopy.

For biofilm samples, the medium was removed and biofilms were gently washed with 3 % BSA in PBS before adding 60 µL BG4 antibody (1: 100) in 3% BSA in PBS and incubating for 90 min with gentle shaking at 50 rpm at room temperature. Without washing or decanting, 60 µL AB1 (1:100) in 3 % BSA was added and samples were incubated for another 90 min under the same conditions. For the next step, biofilms were gently washed with 3% BSA before the addition of 60 µL of the secondary antibody AB2 (1: 100) in 3 % BSA in PBS and 60 min incubation. Finally, biofilms were washed with 3% BSA in PBS and stained with 10 µg/mL FM-464 solution in 100 mM NaCl to visualize the bacteria.

Confocal laser scanning microscopy was performed using a LSM700 microscope (Carl Zeiss) equipped with a 63x objective (Plan-Apochromat, NA1.4 oil immersion objective). We used the bright field transmission to see unstained bacteria, 488 nm excitation and >660 nm emission to visualize FM4-64 stained bacteria, 488 nm excitation and 520 – 600 nm emission for the Atto-488 conjugated BG4 antibody, and 405 excitation and 410-477 nm emission for the AlexaFlour 405-conjugated AB2 antibody.

### Establishment of an experimental model with high or low amounts of G4-DNA

To investigate the contribution of G4-DNA to EET, we first established an experimental model in which bacteria contained many or few G4/hemin complexes associated with the bacterial surface. We hypothesized that G4-DNA would associate with extracellular polysaccharides and therefore acquired a strain in which polysaccharide production could be controlled via an inducible promoter. *S. epidermidis* 1585 pTX*icaADBC* was grown overnight in BHI/NaCl supplemented with tetracycline (20 µg/ml), and xylose (2% w/v) to induce production of poly-n-acetyl glucosamine (PNAG). Bacteria grown in the absence of xylose were used as a polysaccharide-deficient control.

To prepare the experimental model, overnight culture of bacteria were collected in 1 mL broth and centrifuged (5 min at 906 × *g*), washed, and resuspended in 100 mM KCl to an optical density at 600 nm (OD_600_) of 0.5. Annealed G4-DNA (5 µM) and/or hemin (5 µM) was added (5 µM hemin best supported hemin tolerance by *S. epidermidis*), and samples were vortexed (1 min), incubated at room temperature for 1 h in the dark, centrifuged (2 min at 906 × *g*), and resuspended in 100 mM KCl to wash away unbound G4-DNA or hemin. Surface-bound G4- DNA was visualized by fluorescence immunolabeling as described above.

Annealed c-Myc1 and c-Myc3 G4-DNA sequences were adsorbed on PNAG producing *S. epidermidis* 1585 pTX*icaADBC* separately and immunolabelled for comparison. After confirming that *S. epidermidis* 1585 pTX*icaADBC* could adsorb G4-DNA to the bacterial surface when PNAG production was induced, this strain was used as a model organism with high or low amount of G4-DNA structures in the subsequent experiments. All subsequent experiments were done with the G4-DNA sequence with highest binding.

Prior to the experimental model, concentrations of G4-DNA/hemin concentration tolerable to planktonic cells were determined and used for subsequent experiments with the *S. epidermidis* 1585 pTX*icaADBC* strain. BHI/NaCl broths supplemented with tetracycline (20 µg/ml), with and without xylose (2 % w/v) were prepared to test the effect of PNAG production by planktonic cells on hemin tolerance. Overnight cultures were collected in 1 mL broths, centrifuged (5 min at 906 *× g*) and pellets were resuspended in same medium broths to an optical density (OD_600_) of 0.01 in 2 mL volumes before transferring to sterile 48 well plates (Sarstedt AG & Co). Different hemin concentrations (0, 5, 40 and 200 µM) +/- 5 µM, annealed c-Myc3 were then added. Unlike hemin, G4-DNA was not considered toxic, and concentration used (5 µM) in this experiment was kept constant. Experiments were carried out in triplicates and kept in the dark. Microtitre plates with culture were incubated for 24 h with shaking (100 rpm) at 37 °C. After incubation, 0.1 mL of each of the cultures were serially diluted 10-fold in sterile deionized water. Dilution 10^-7^ was plated onto BHI agar by transferring 0.1 mL on the agar. Inoculum were evenly spread on the agar using sterile cell spreader. Agar plates were incubated at 37 °C for 24 h. Colonies were counted after incubation and CFU/mL was determined. Logarithmic values of the CFU/mL were calculated and recorded.

### Electroanalysis of planktonic bacteria adsorbed to the electrode

*S. epidermidis* 1585 pTX*icaADBC* was grown overnight in BHI/NaCl supplemented with tetracycline (20 µg/ml), and +/- xylose (2% w/v) to induce polysaccharide production. Bacteria were washed by centrifugation (5 min at 1118 × *g*) and adjusted to OD_600_ 0.5 in 100 mM KCl with G4-DNA and/or hemin treatments as described above. For comparison, electroanalyses with planktonic *S. epidermidis* 1457 and *S. epidermidis* 1457 Δ*atlE* were also carried out on overnight cultures grown in BHI/NaCl, washed and suspended in 100 mM KCl to OD_600_ 0.5 with G4-DNA + hemin treatment as described above.

The SPEs (Metrohm DropSens DRP-C110, Metrohm Spain) had a 4 mm diameter transparent Indium Tin Oxide (ITO) working electrode (WE) with surface area of 0.126 cm^2^, a graphite counter electrode, and Ag pseudo-reference electrode. Prior to use, SPEs were sterilized in 70% v/v ethanol, washed thrice in sterile deionized water, and air dried. Bacterial suspensions (100 μL) were then deposited on screen printed electrodes (SPEs) and left to adsorb for 15 min. During deposition, cells were made to cover the whole surface area of working, reference and counter electrode. Excess cells were gently rinsed with 100 mM KCl and SPEs were subsequently connected to a computer-controlled multichannel PalmSens 4 potentiostat (PalmSens® Netherlands) equipped with an EmStatMUX8-R2 multiplexer.

All electrochemistry data were collected and analysed using the PalmSens PsTrace 5.9 software. Cyclic voltammetry (CV) was run between -0.8V and 0.8V with a step size of 0.01 V and a scan rate of 0.05 V/s for 3 scan cycles, and the last scan was selected per sample. Samples were run in triplicates and the averages were plotted. Differential pulse voltammetry (DPV) was carried out by first pretreating at 0.2 V for 2 min before running with -0.8 V as start potential and 0.8 V as end potential, with scan rate of 0.05 V/s, step size 0.01 V, pulse time 0.02 s and current range from 10 nA to 1mA. The average scans of three biological replicates were recorded for each voltammetric experiment.

Chronoamperometry (CA) was also carried out using 0.4 V poised potentials for 300 s after 200 s pre-treatment and delay time to allow the bacteria to acclimatize to the poised potential. Values of total charge generated were also recorded after the CA measurements. Multi-step chronamperometry (MSCA) was carried out by switching between poised potentials 0.4 V and -0.4V for 30 s with 3 s between switches.

At the end of the experiment, the adsorbed bacteria and the presence of G4-DNA structures were visualized by immunolabeling and microscopy as described above. Comparative CV analyses was carried out on c-Myc1 and c-Myc3 G4DNA treated planktonic cells of polysaccharide producing *S. epidermidis* 1585 pTX*icaADBC* in the presence of hemin. G4- DNA with higher electroactivity was selected for further analyses.

### Blocking of electron transfer with G4-DNA-binding antibodies

We sought to validate the contribution of G4-DNA/hemin to electroactivity, as hemin could potentially also associate with other biomolecules on the bacterial surface. We therefore used a G4-DNA specific antibody to block the electron flow from G4-DNA/hemin on bacteria to the electrode. Overnight cultures of *S. epidermidis* 1585 pTX*icaADBC* were prepared, suspended in 100 mM KCl, treated with G4-DNA and hemin, and deposited on SPEs as described above. However, before depositing bacteria, SPEs were treated with 60 µL BG4 antibody (1: 100 in 3% BSA in PBS) or a non-specific antibody (anti-mouse antibody - goat monoclonal IgG H + L, XF346503 Invitrogen®), and allowed to dry for 30 min. As controls SPEs were treated with BG4 antibody alone or G4-DNA (5 µM). CA was carried out using 0.4 V poised potentials as described above.

### Electroanalysis of *S. epidermidis* biofilms grown directly on screen printed electrodes

We first aimed to use a wildtype *S. epidermidis* (strain 1457) and an eDNA-negative mutant (*S. epidermidis* 1457 Δ*atlE*) to determine if *S. epidermidis* biofilms are electroactive, and if this activity correlates with the presence of eDNA. We used *S. epidermidis* 1457 for this purpose because previous publications had documented formation of G4-DNA in the biofilm eDNA network formed by this strain.

Overnight cultures grown in TSB/NaCl were centrifuged (2 min at 5000 rpm) and transferred to TSB/NaCl with 5 µM hemin, adjusted to OD_600_ of 0.05, and 1.5 mL was then transferred to sterile 2 mL polystyrene cuvettes (Sarstedt AG & Co). Sterile SPEs were inserted into the solution (the connective end of SPEs was not submerged) and incubated at 150 rpm for 48 h before analysis by CV and DPV to determine the electrochemical peaks as described above. The average scans of three biological replicates were recorded for each voltammetric experiment. As a control experiment to separate the activity of biofilms from planktonic bacteria, planktonic cultures of both strains were grown for 48 h in TSB/NaCl + 5 µM hemin, and sterile SPEs were inserted into the culture for DPV analyses of the planktonic culture.

*S. epidermidis* 1457 may not naturally produce a large and controllable amount of G4-DNA structures in the biofilm. To better understand how G4-DNA can contribute to EET in biofilms, we therefore used our experimental model *S. epidermidis* 1585 +/- PNAG production and +/- addidion of G4-DNA oligos to study EET in biofilms with high or low amounts of G4-DNA structures in the extracellular matrix.

*S. epidermidis* 1585 pTX*icaADBC* biofilms were grown in 1 mL anaerobic custom-made containers mounted directly on the SPEs. A 10 mm diameter glass vial with butyl rubber cap was cut to remove the glass bottom. The top was then attached to the SPE using non-conductive glue as described in Han et al. ^53^ with slight modifications. The container was sterilised with 70% ethanol and UV light for 15 min. BHI/NaCl (0.5 mL) with 2% xylose and/or hemin (5 µM) and G4-DNA (5 µM) was added and sparged with N_2_ gas for 2 min using sterile needles to remove oxygen before inoculation with OD_600_ 0.1 *S. epidermidis* 1585 pTX*icaADBC* overnight cultures grown in BHI/NaCl + 20 µg/mL tetracycline + 2% xylose. Control experiments were designed to determine the function of iron containied in hemin in redox transfer. We grew and treated *S. epidermidis* 1585 pTX*icaADBC* in similar conditions stated above but replaced hemin with 5 µM protoporphyrin IX (PPIX) which had similar structure with hemin, but lacked iron. Electrodes were poised at 0.4 V and incubated at 37°C. The potential induced biofilm formation on the electrode, using it as an electron acceptor in the absence of oxygen. Time-based electroanalyses (CA, DPV and charge measurements) were carried out on the growing biofilms for up to 120 h using parameters described above. Immunolabelling for time-based experiments, involved the use of different electrode set-ups for each time considered and immunolabeling and microscopy were subsequently conducted as described above.

### Detection of G4-DNA/hemin peroxidase activity

The G4-DNA/hemin complex is also a DNAzyme with peroxidase-like activity. It could therefore also contribute to EET by transferring electrons to alternative electron acceptors, such as hydrogen peroxide (H_2_O_2_). We validated the peroxidase activity of the G4-DNA/hemin adsorbed to *S. epidermidis* 1585 pTX*icaADBC* using tyramide signal amplification (TSA). This method is commonly used in immunolabelling with horseradish peroxidase (HRP)-conjugated antibody. In our assay, we visualized the location of peroxidase activity stemming from the G4- DNA/hemin DNAzyme before proceeding to test if the DNAzyme can catalyse the transfer of electrons from bacteria to H_2_O_2_.

Overnight cultures of *S. epidermidis* 1585 pTX*icaADBC* were grown in BHI/NaCl supplemented with tetracycline (20 µg/ml), and xylose (2% w/v) and contained a combination of planktonic bacteria and larger aggregates. Twenty µL culture was transferred to an Eppendorf tube with a cut-off pipette tip to avoid shearing of the aggregates. Aggregates were suspended in 200 µL modified MES buffer adjusted to pH 6.5. Samples were supplemented with G4-DNA (5 µM) and/or hemin (5 µM) and vortexed for (30 s) before transferring 90 µL to a coverslip with an attached Gene Frame (Thermo Scientific, AB0577) and incubating for 30 min (RT, 50 rpm shaking) to let the bacteria and aggregates adsorb before removing the liquid and replacing it with 90 µL tyramide reagent containing tyramide conjugated with Alexa Fluor 488 (1:100) (Invitrogen, B40953), 2 mM ATP (Thermo Scientific, R0441), and 0.1% hydrogen peroxide (Sigma, 216763) in modified MES buffer. Samples were then incubated for 90 min (RT, 50 rpm). Samples were washed with modified MES buffer, and stained for 30 min (RT, 50 rpm), with 90 µL of 20 µM SYTO^TM^ 60 (Invitrogen, S11342) prepared as 100 x aliquots in sterile deionized water. Samples were then mounted with Prolong^TM^ Glass Antifade Mountant (Invitrogen, P3698) after a gentle wash before CLSM imaging as described above. Tyramide-AlexaFLour-488 was excited at 488 nm and emission detected from <550 nm. SYTO^TM^ 60 was excited at 647 nm and emission detected at 650-700 nm. Both lasers were scanned in the same track with a beam splitter at 560 nm. Planktonic cells prepared as described above but not stained with Syto 60, were also used. Bright field images showed the planktonic cells during tyramide assay.

### Electrochemical detection of residual hydrogen peroxide after degradation

After confirming peroxidase activity of the G4-DNA/hemin complex, we proceeded to determine if bacteria could transfer electrons from their metabolism to H_2_O_2_ via G4- DNA/hemin. We measured the capacity for bacteria to remove H_2_O_2_, comparing *S. epidermidis* 1585 pTX*icaADBC* in the presence or absence of hemin, G4-DNA or both.

Overnight cultures of *S. epidermidis* 1585 pTX*icaADBC* were prepared as described for the TSA assay, transferred to an Eppendorf tube, diluted to OD_600_ of 1 in fresh media and supplied with G4-DNA (5 µM) and/or hemin (5 µM). 200 µL culture was centrifuged (at 1118 × *g* for 2 min) and washed in same volume of 0.1 M KCl. Cells were resuspended in 200 µL modified MES buffer in 500 µL tubes and sparged with N_2_ gas for 30 s before addition of 0.1% hydrogen peroxide and incubating at 37°C with gentle shaking (40 rpm) for 5 min. After incubation, cultures were centrifuged at 1118 × *g* for 2 min, and the supernatant was collected. The supernatant (50 µL) was dropcast on sterile SPEs connected to potentiostat and cyclic voltammetry was immediately run from 0.8 V to -0.8 V at 50 mV/s to detect H_2_O_2_. Buffer containing undegraded 0.1% H_2_O_2_ served as the first control. G4-DNA (5 µM) mixed with hemin (5 µM) in 200 µL modified MES buffer containing 0.1% H_2_O_2_ were incubated anaerobically and used as a second control.

## EXTENDED DATA

**Extended Data Fig. 1:**
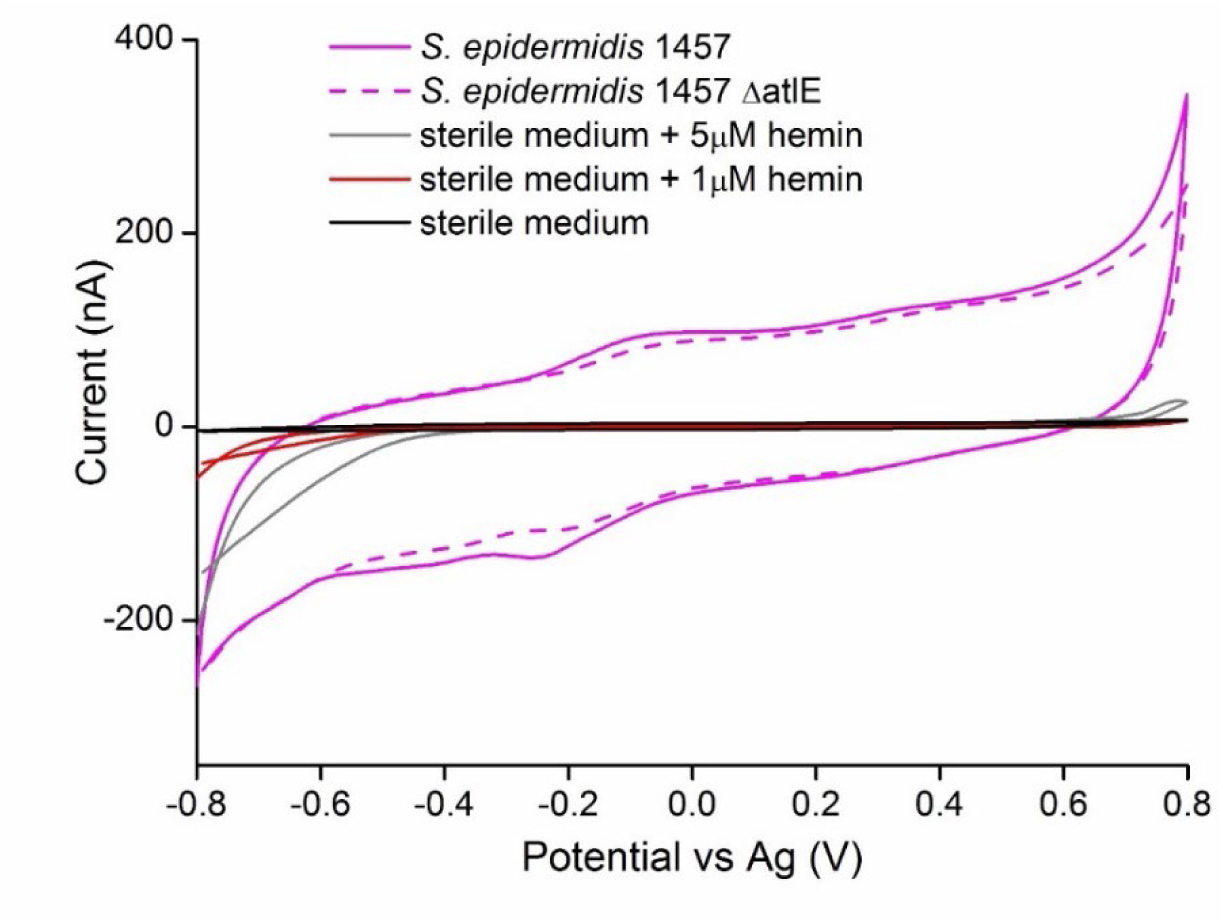
Cyclic Voltammogram (CV) of biofilms formed by *S. epidermidis* 1457 (eDNA producing) and *S. epidermidis* 1457 Δ*atlE* (autolysin deficient and low eDNA producing) when grown with 5 µM hemin for 48 h. Sterile TSB/0.2 M NaCl medium and Sterile medium (+1 µM and +5 µM hemin) served as controls.

**Extended Data Fig. 2:**
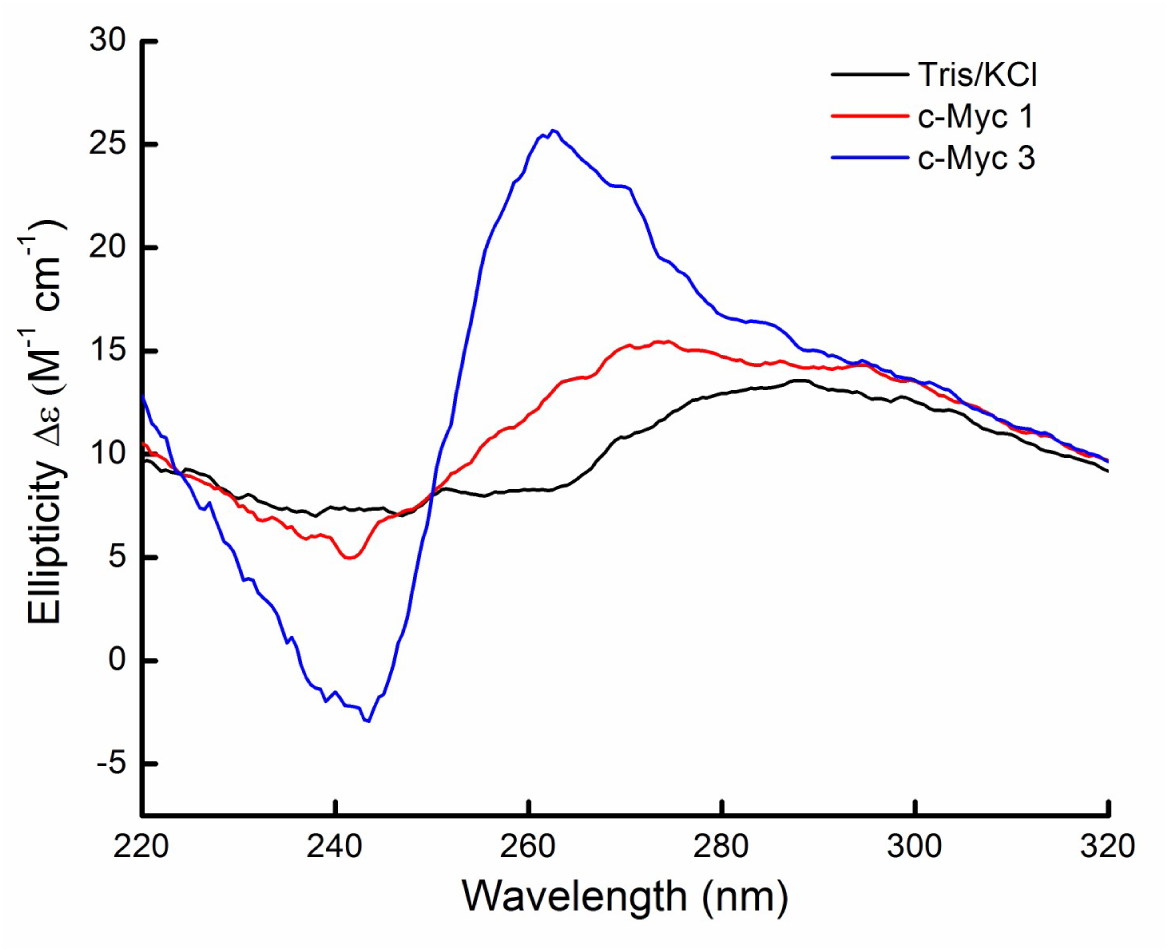
Circular dichroism confirms the structure of G4-DNA oligonucleotides

**Extended Data Fig. 3:**
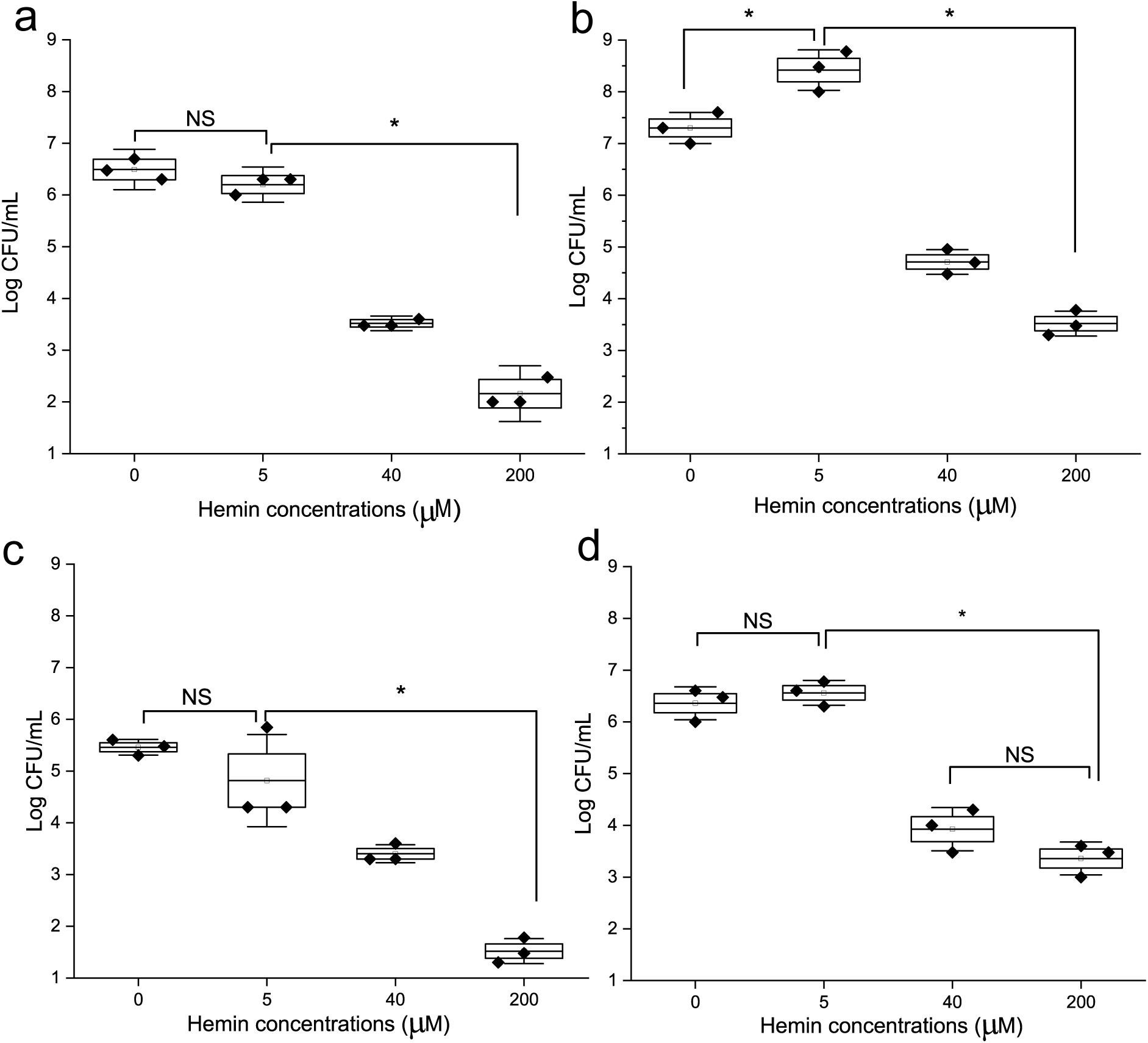
Cell viability and tolerance tests of planktonic cultures of *S. epidermidis* grown in varying concentrations of hemin with or without PNAG induction and G4-DNA (c-Myc3, 5 µM) addition (a) PNAG -, G4-DNA + (b) PNAG +,G4-DNA + (c) PNAG -, G4-DNA - (d) PNAG +, G4-DNA –. ∗ represents statistical significance (p < 0.05), ANOVA and Tukey’s test, NS= Not significant.

**Extended Data Fig. 4:**
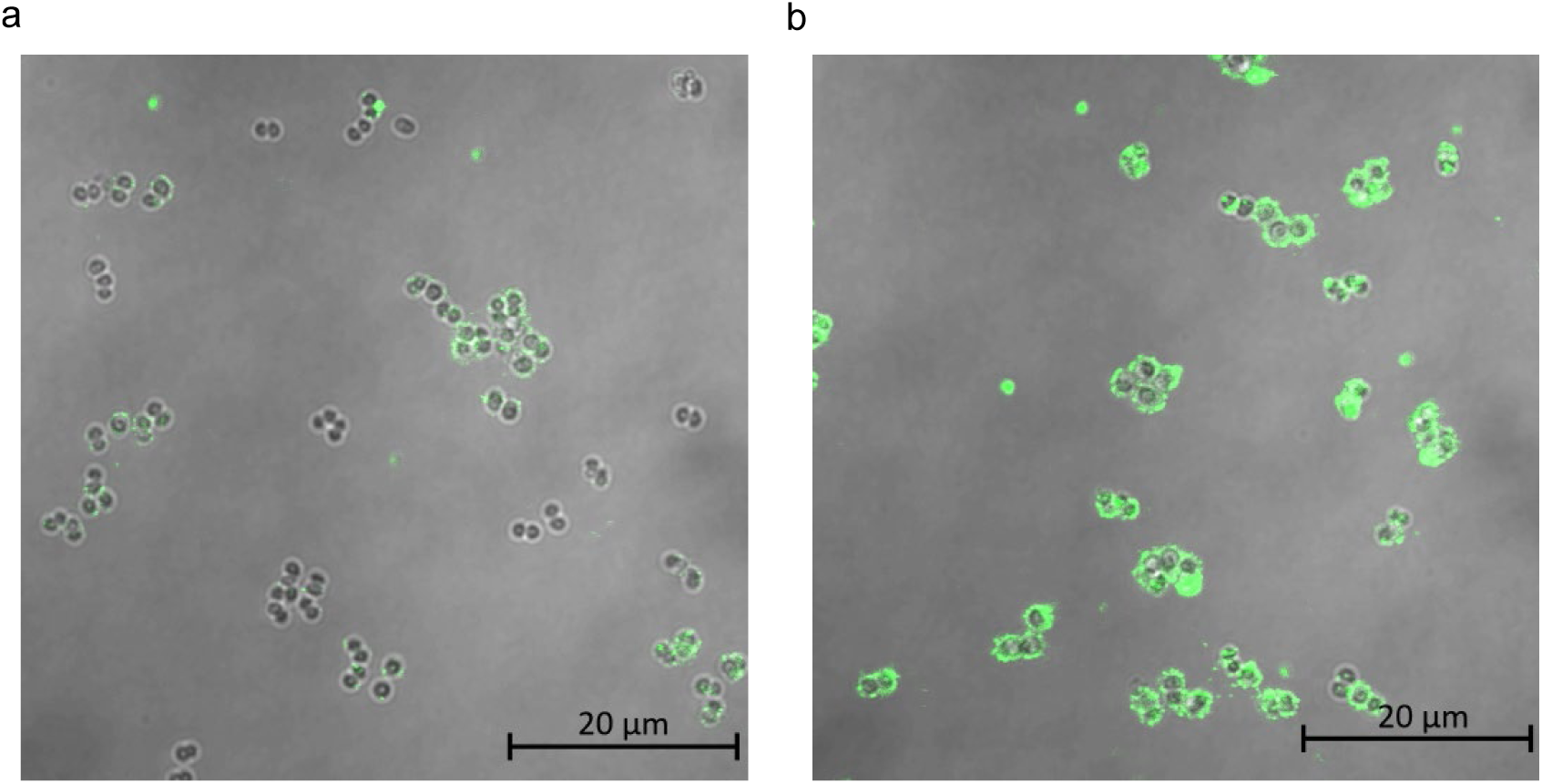
Adsorption of c-Myc1 (A) and c-Myc3 (B) to the surface of *S. epidermidis* with induced PNAG production. More G4-DNA was detected on the bacterial surface when using c-Myc3, and this oligo was therefore used in our subsequent experiments.

**Extended Data Fig. 5:**
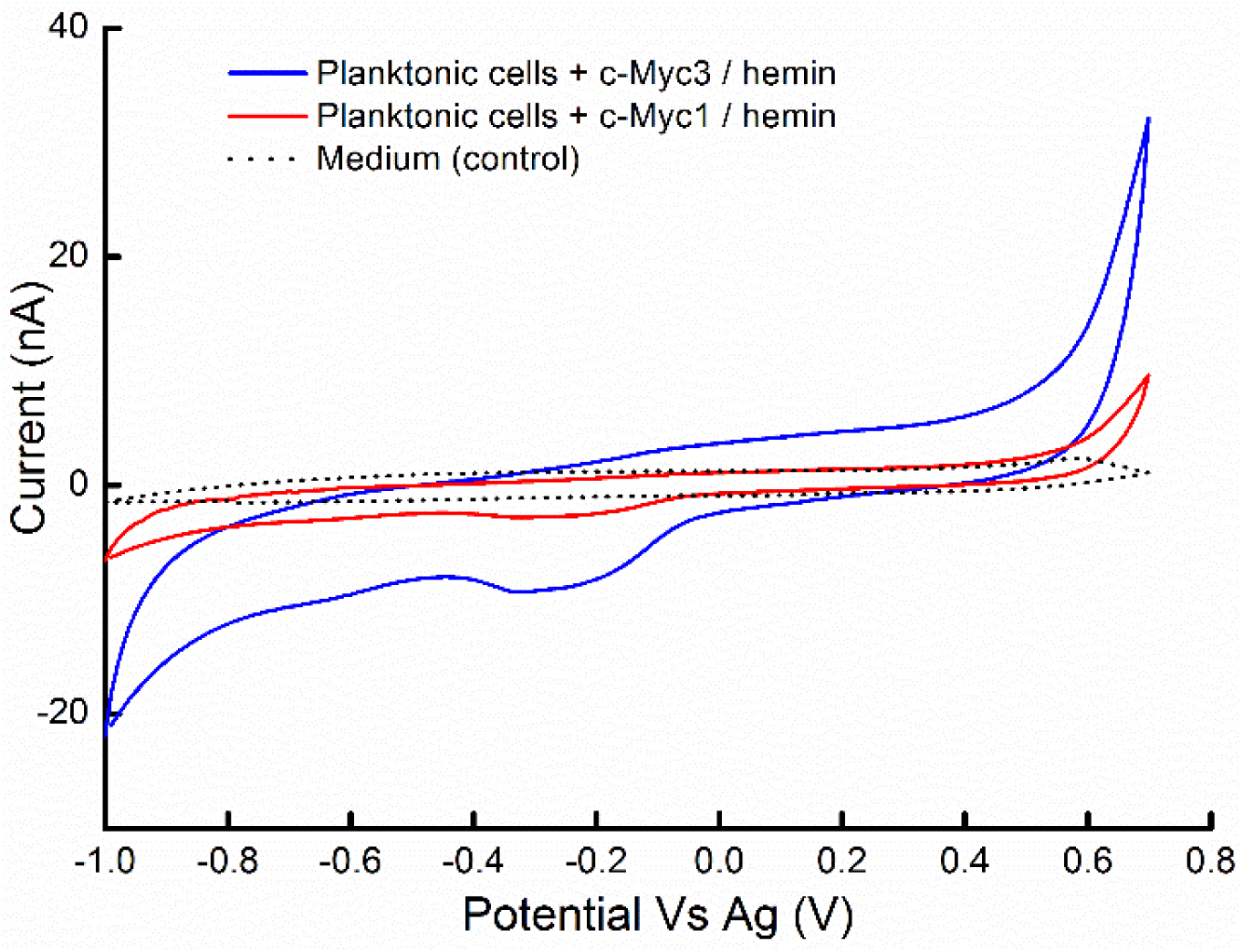
Cyclic Voltammogram (CV) shows that bacteria with c-Myc3 are more electroactive. Planktonic *S. epidermidis* were grown in TSB/NaCl with xylose (for PNAG production) and treated with 5 µM G4-DNA and 5 µM hemin before adsorption to the electrode and measurement of CV.

**Extended Data Fig. 6:**
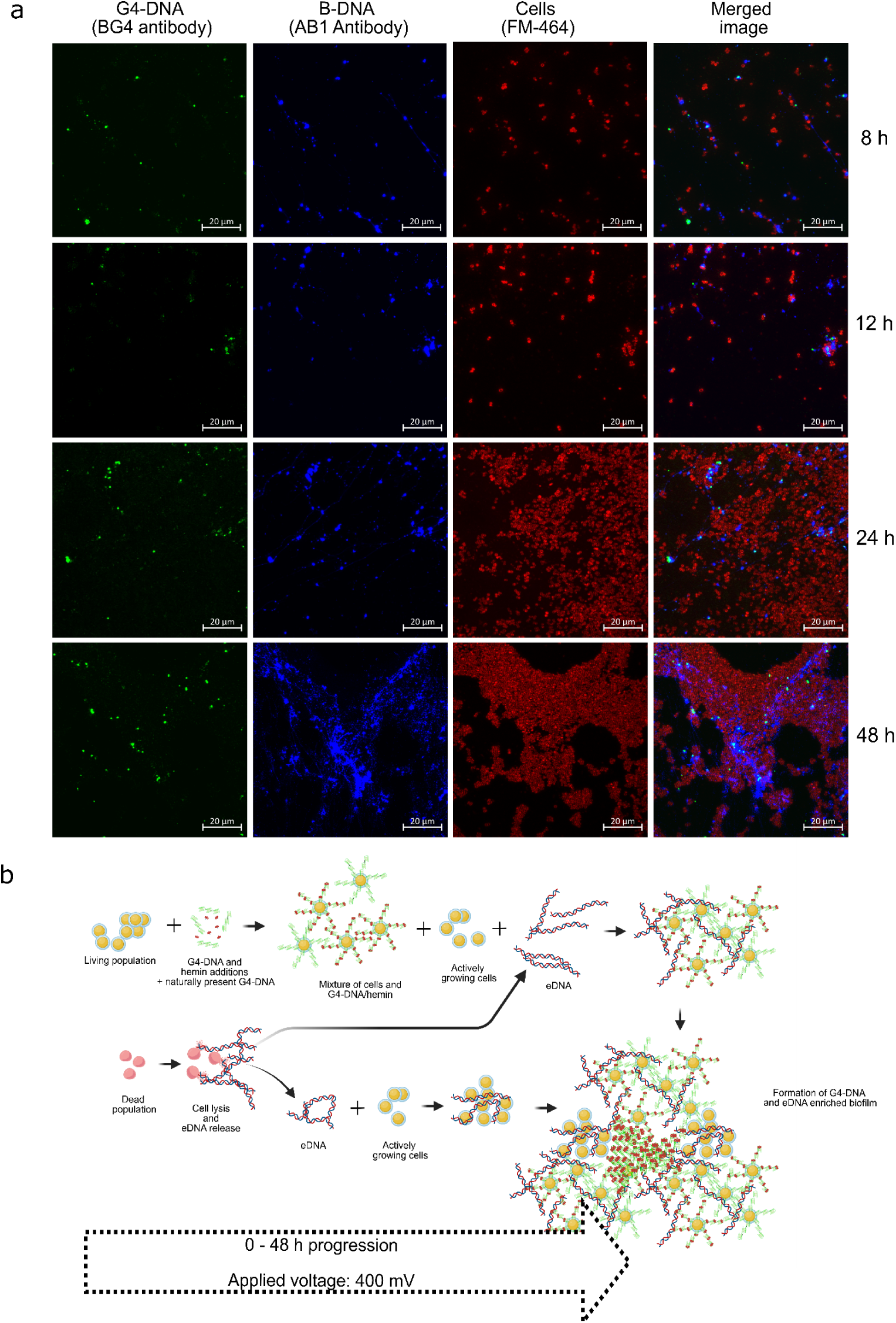
Externally added G4-DNA are gradually recruited over time by eDNA formed in the biofilm matrix. Time-based recruitment of free-floating G4-DNA for formation of enriched biofilm with G4-DNA/hemin complex under electrical influence of 0.4V poised potential (a) Microscopic observation of biofilm formation from 8h to 48h showing gradual G4-DNA incorporated over time into the biofilm under electrical influence (b) Proposed scheme for G4-DNA enriched biofilm formation over 48h. We propose that initial eDNA (presented as B-DNA) start to appear due to lysis of some cells from the starting inoculum after first exposure to applied electric potential. Gradual incorporation of G4-DNA into electroactive biomass leads to interactions between B-DNA and G4-DNA, as well as G4- DNA and cells. (b) is created in BioRender. Ajunwa, O. (2024) https://BioRender.com/h06f025

**Extended Data Fig. 7:**
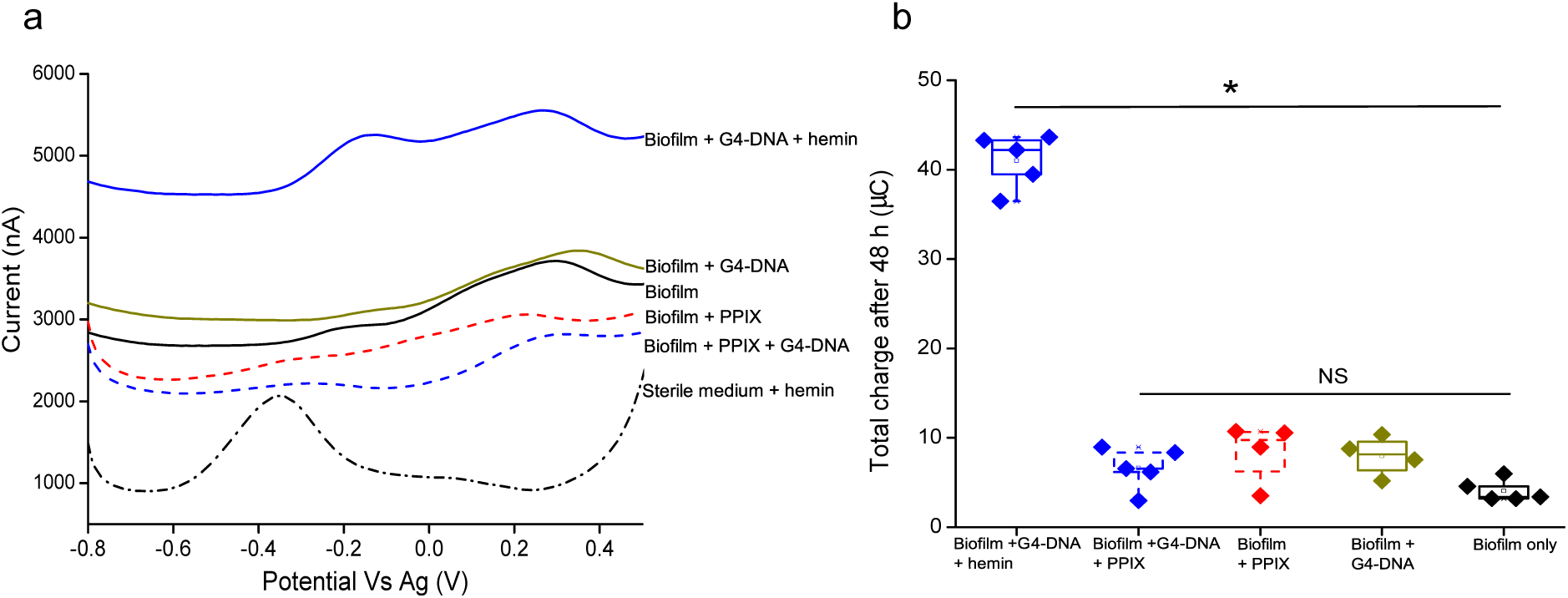
Biofilms grown with G4-DNA and hemin facilitate EET in an iron dependent fashion. Comparative electroactivity of 48 h old biofilms of *S. epidermidis* with induced PNAG production. Biofilms were supplemented with G4-DNA and either hemin or the iron deficient PPIX. (A) DPV shows PPIX treated biofilms lack the characteristic negative potential peak (-0.3 V to 0.1 V) associated with G4-DNA/hemin interactions (B) Total electric charge generated from biofilms also confirm the requirement of iron porphyrin to facilitate charge transfer (∗ represents significant difference (p < 0.05) based on ANOVA and Tukey’s test, NS: Not significant).

**Extended Data Fig. 8.**
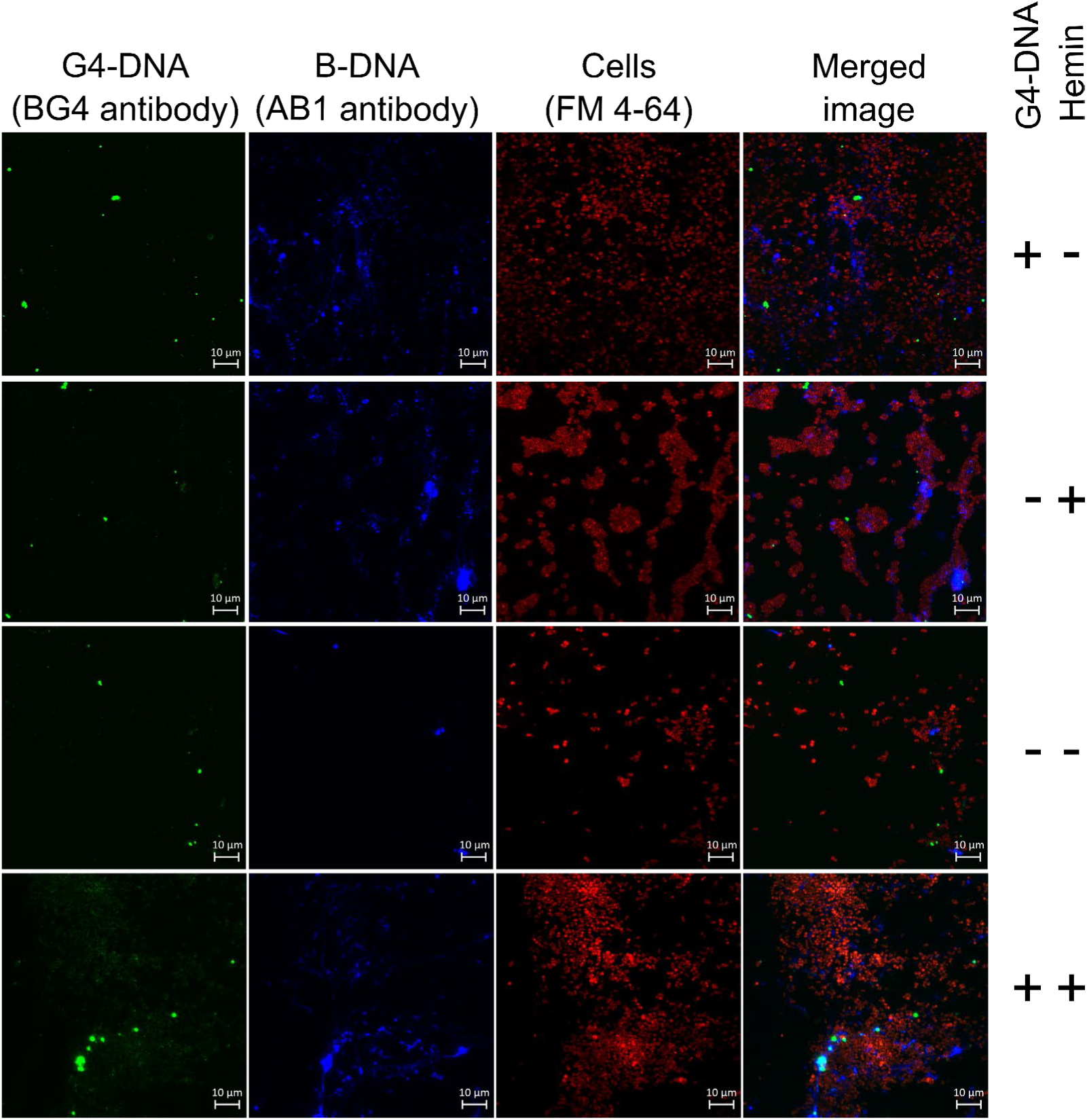
Representative microscopy images of *S. epidermidis* 1585 pTX*ica* biofilms formed on electrodes after 48 h with different treatments of G4-DNA (+/-) and hemin (+/-). Samples with both G4-DNA and hemin forms nodular structures of G4-DNA that associate with B-DNA in the biofilm.

**Extended Data Fig. 9.**
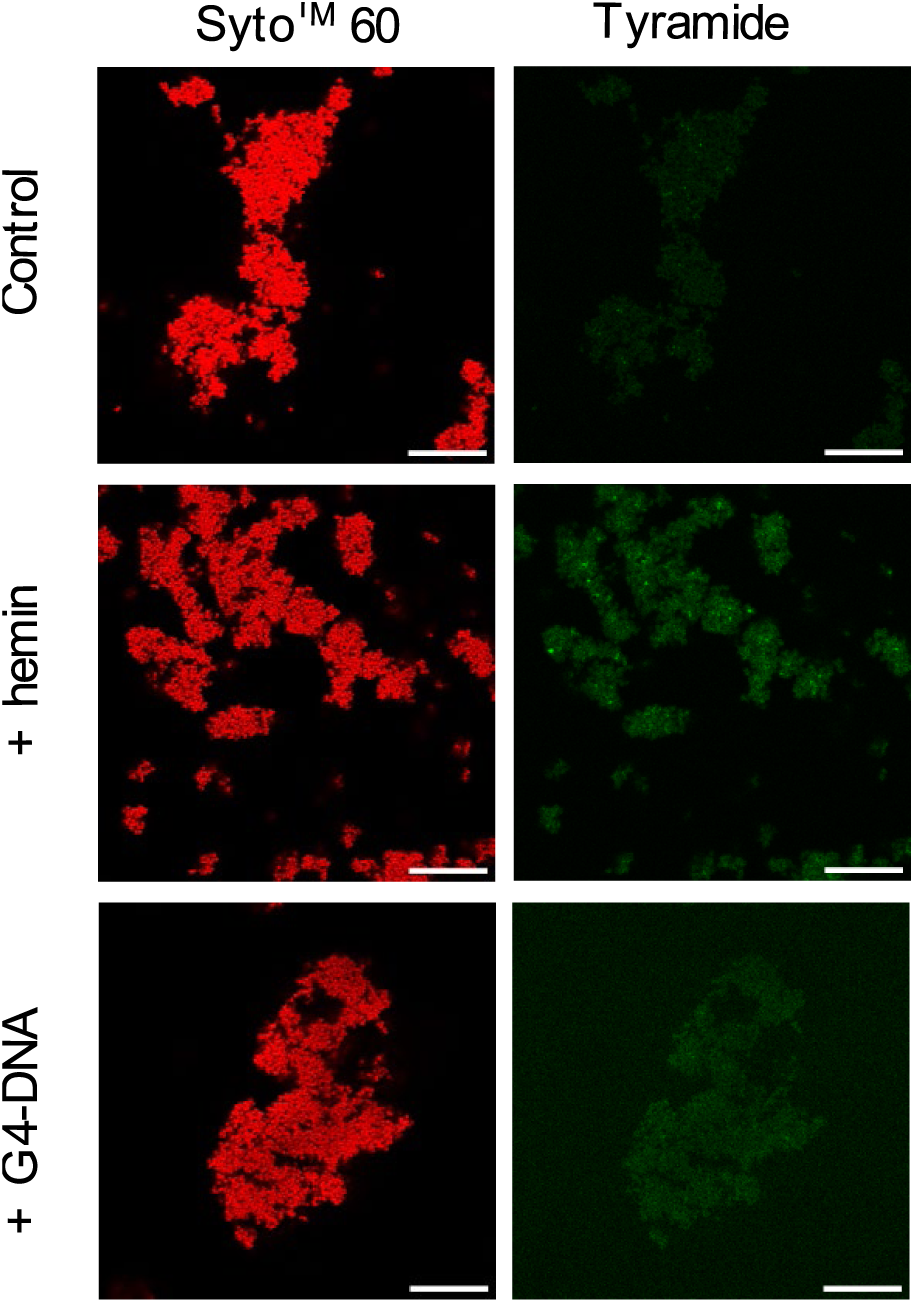
Lack of G4-DNA/hemin on the bacteria results in low peroxidase-like activity. Tyramide signal amplification shows low signals around the cells after reacting with tyramide and hydrogen peroxide. CLSM of bacterial aggregates stained by Syto 60 (red). Peroxidase activity showing minimal fluorescence at sites of peroxidase activity. Scale bar = 20 µm.

